# Assembly of functional diversity in an oceanic island flora

**DOI:** 10.1101/2022.03.04.482684

**Authors:** Martha Paola Barajas-Barbosa, Dylan Craven, Patrick Weigelt, Pierre Denelle, Rüdiger Otto, Sandra Díaz, Jonathan Price, José María Fernández-Palacios, Holger Kreft

## Abstract

Oceanic island floras are well-known for their morphological peculiarities and exhibit striking examples of trait evolution^1,2^. These morphological shifts are commonly attributed to insularity and thought to be shaped by biogeographical processes and evolutionary histories of oceanic islands^1,3^. However, the mechanisms through which biogeography and evolution have shaped the distribution and diversity of plant functional traits remain unclear. Here, we describe the functional trait space of an oceanic island flora (Tenerife, Canary Islands, Spain) using extensive field and laboratory measurements, and relate it to global trade-offs in ecological strategies. We find that the island trait space is concentrated around a functional hotspot dominated by shrubs with a conservative life-history strategy. By dividing the island flora into species groups with distinct biogeographical distributions and diversification histories, our results reveal that long-distance dispersal, and the interplay between inter-island dispersal and archipelago-level speciation processes drive functional divergence and expand trait space. Conversely, speciation via cladogenesis has overall led to functional convergence, densely packing trait space around shrubbiness. Our approach combines ecology, biogeography and evolution and opens avenues for new trait-based insights into how dispersal and speciation shape the assembly of native island floras.

## Main text

Oceanic islands have attracted enormous interest in biogeography^4,5^ and serve as natural laboratories to study the assembly of floras and faunas^6,7^. Empirical tests of fundamental concepts in evolution and ecology using islands as model systems^2,8^ assume that the results can be generalised to non-island contexts. Yet, a long-standing paradigm in island biogeography centres on the notion that isolation, as well as geographic and environmental factors linked to the ontogeny of oceanic islands, lead to evolutionarily unique^9,10^ and functionally distinct biotas^11^. These assumptions raise the question of how functionally distinct oceanic islands can be compared to other systems.

In contrast to neutral theories, such as the theory of island biogeography^8^, where all arriving species have the same chance to colonise an island irrespective of their traits, a trait-based perspective leverages differences in ecological strategies among species determined by functional traits to disentangle the processes and identify the mechanisms that have shaped island biotas^3,12^. In functional ecology and functional island biogeography^12,13^, functional differences among species are quantified using morphological and physiological characteristics that impact how plants respond to environmental factors, affect other trophic levels, and influence ecosystem properties^14,15^. Trait syndromes and trade-offs among individual traits reflect fundamental ecological strategies that structure plant life from individuals to communities. This includes the conservative-acquisitive continuum captured by the leaf economics spectrum^16,17^ and the size continuum (stature of whole plants and seed mass), which together, define essential dimensions of functional diversity in vascular plants (e.g., the global spectrum of plant form and function, *sensu*^18^). Functional traits are also used to estimate functional diversity^19^, which quantifies the diversity and distribution of ecological strategies in a community^20^ and may help identifying the factors that shape an assemblage, such as abiotic conditions^21,22^ or dispersal filters^10^. Ecological, biogeographical, and evolutionary forces may either expand or constrain trait space, thereby driving functional divergence or convergence^23^. However, shortfalls in trait and distribution data^24,25^ usually restrict the geographic extent, spatial grain, and taxonomic coverage of such studies, leading to an incomplete and potentially biased understanding of the factors that underpin a complete flora.

Crossing large expanses of ocean and adapting to island environments represents a considerable challenge for plants, likely filtering species with trait values that enhance dispersal and establishment^10^ (but see^26^). Moreover, the highly heterogeneous environments^27^ of oceanic islands create ecological opportunities (i.e., empty niche spaces)^9^ that should promote divergence of new trait combinations. Biogeographical processes, such as long-distance dispersal, are thus expected to strongly constrain an island trait space, leading to functional convergence, where species share similar trait combinations. In contrast, evolutionary events, such as speciation via lineage splitting (cladogenesis^28^) or via gradual evolution of species (i.e., anagenesis^29^) are expected to expand trait space^30^ by generating novel trait combinations that facilitate the occupation of untapped trait space. The Hawaiian silverswords and the Macaronesian *Echium* alliance are iconic examples of adaptive radiations involving dramatic morphological shifts^31,32^, yet, the relative importance of biogeographical and evolutionary processes in shaping trait diversity and ecological strategies on oceanic islands remains elusive.

Here, we study the flora of Tenerife (Canary Islands, Spain), an oceanic island located in Macaronesian meta-archipelago and an ideal natural laboratory^33^ to test how biogeography and evolution affect island functional diversity. Tenerife exhibits stunning environmental gradients from arid coastal succulent scrub vegetation, to humid laurel forests and alpine vegetation^34^ and contains spectacular examples of insular radiations^33^. Tenerife’s flora is well-studied and comprises 436 native seed plants^35^. We measured eight plant functional traits of 80% of the native seed plants (Fig. 1 and Fig. 2a) to explore matches and mismatches in trait syndromes between Tenerife’s native flora and the global spectrum in plant form and function^18^ (Fig. 1). We evaluate how biogeographical and evolutionary processes have shaped the functional diversity of Tenerife’s flora by assessing the contribution of five species groups, which are associated with different dispersal and speciation mechanisms, to the overall island trait space (Fig. 2b-f): (i) non-endemic native species, which represent species that colonised the Canaries via long-distance dispersal without undergoing subsequent speciation^36^, (ii) Macaronesian endemics, which signal ‘relictualization’, a process that results from the survival of species on islands after the extinction of mainland populations, as well as speciation^37,38^, (iii) Canary Islands endemics, which represent both speciation and inter-island dispersal across the archipelago, (iv) Tenerife endemics, which represent *in-situ* speciation and intra-island dispersal. As a fifth group, we also considered (v) cladogenetic species, which is composed of immigrant lineages that diversified within the Macaronesian meta-archipelago.

**Figure 1.**
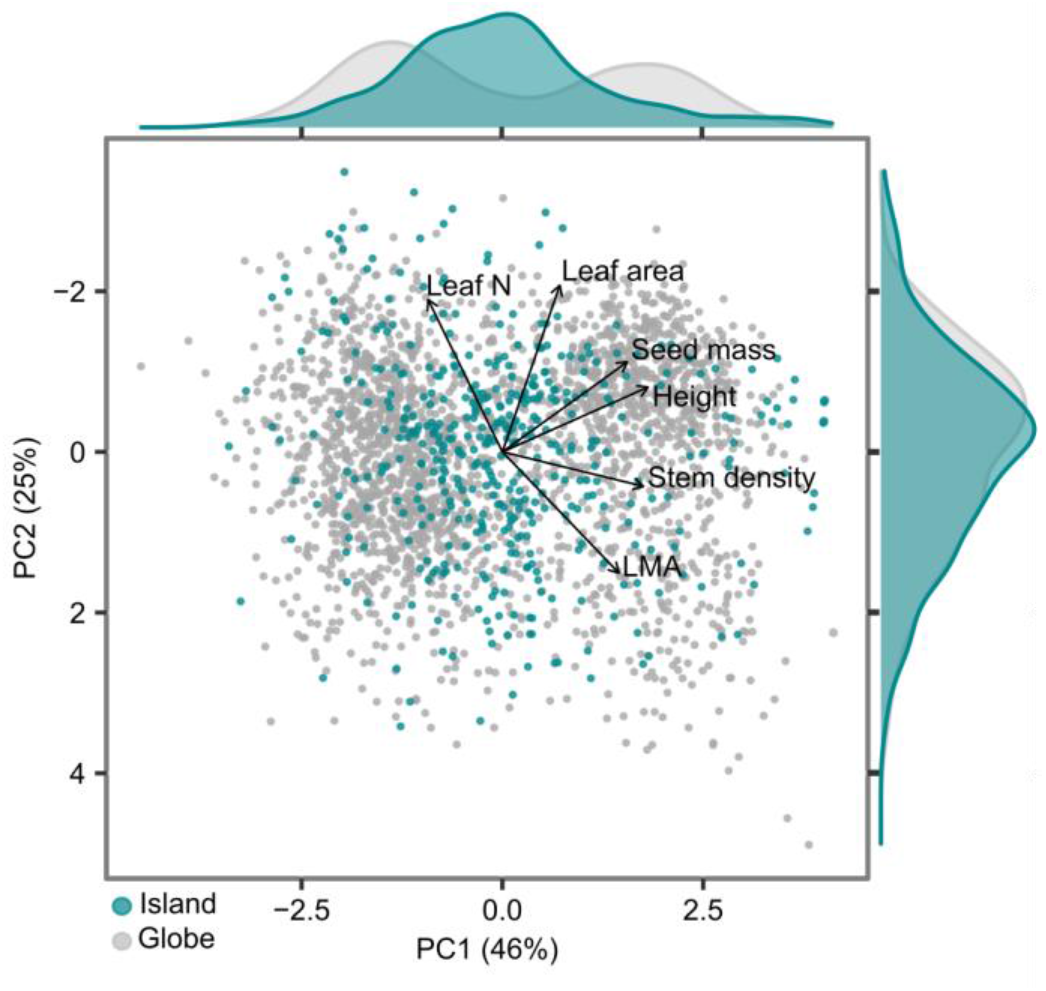
Trait space of the native flora of Tenerife is subject to similar constraints as the global trait space, but is biased towards medium-statured species with intermediate trait values. Trait space of Tenerife using 436 native seed plant species (turquoise dots) in relation to the global plant trait space known as the global spectrum of plant form and function (2199 species, grey dots, Díaz et al. 2016). Projections of the first two dimensions of variation from a principal component analysis (PCA; for comparability we inverted the y axis, i.e., PC2) of six plant functional traits: leaf area (mm^2^), leaf mass per area (LMA; g/m^2^), leaf Nitrogen content (Leaf N; mg/g), maximum plant height (Height; m), stem specific density (Stem density; mg/mm^3^) and seed mass (mg). Density distributions of the first and second dimensions of island and global trait spaces are displayed in the upper and right side of the trait spaces. More detail on the PCA in Extended Data Fig. 1.

**Figure 2.**
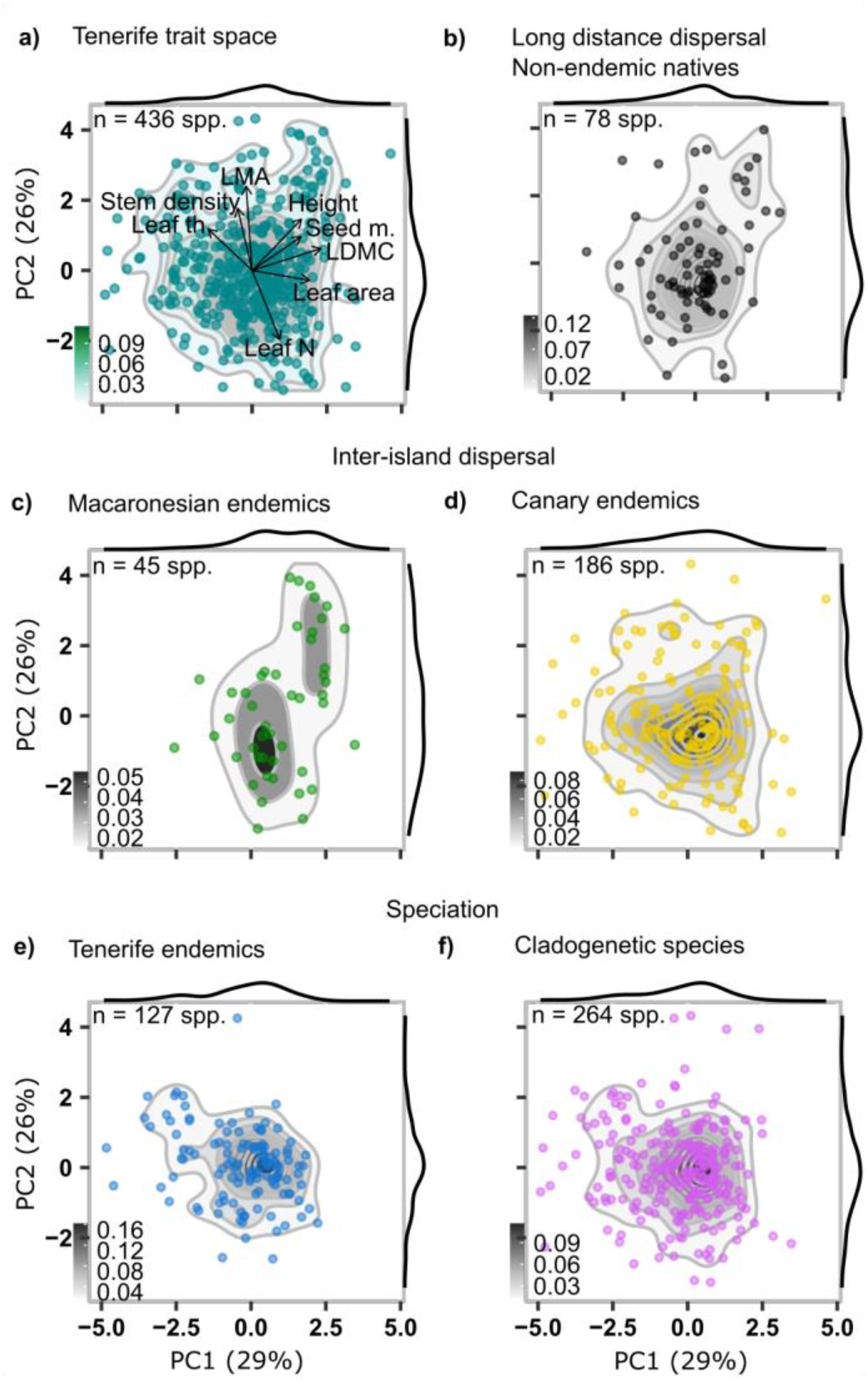
Trait spaces for the native flora of Tenerife(a) and dissected into five distinct species groups (b-f) illustrate how biogeography and evolution shape the functional diversity of an oceanic island flora (n = 436 species). Traits included are leaf dry matter content (LDMC), leaf mass per area (LMA), Leaf N (leaf Nitrogen content), leaf thickness (Leaf th), stem density (SSD), seed mass (Seed m.), and maximum plant height (Height). Projections are of the two first dimensions of variation from the principal component analysis. Contours are built using 2D kernel density estimation. Gradient legends (bottom-left side) correspond to the proportion of data contained in a contour break. More detail on the PCA in Extended Data Fig. 2.

### An oceanic island flora faces similar functional constraints as other floras across the globe

Comparing the variation of traits in the flora of Tenerife with the global spectrum of plant form and function^18^ (Fig. 1 and Extended Data Fig. 1), we find a considerable overlap between the island and the global trait space (Sørensen similarity coefficient = 0.69, based on hypervolume overlap for both trait spaces). This suggests that plants on Tenerife largely experience similar constraints as plants in continental floras. However, the density distribution of Tenerife’s trait space deviates strikingly from the global one along the first trait space dimension (Fig. 1). The majority of island species are located between the two global functional hotspots, i.e., small statured, light-seeded herbaceous plants and tall, heavy-seeded trees. The position of Tenerife’s plants in relation to the global trait space indicates the dominance of shrubs and an underrepresentation of both small herbs and tall trees on the island (Fig. 1 and 2a). Most of Tenerife’s plants have a small to intermediate stature (average plant height = 2 m), moderate stem density (stem specific density average = 0.5 mg mm^−3^), light seeds (seed mass average = 19 mg), and leaves with intermediate size, leaf mass per area (LMA) and nitrogen content (average of leaf area = 3.436 mm^2^, LMA = 93 g m^−2^ and leaf nitrogen = 19 mg g^−1^, respectively) (Extended Data Fig. 3a).

Tenerife’s Mediterranean climate, marked by seasonal summer droughts and high aridity at low and high elevations, favours shrubs^39,40^. Major aridification across the Canary Islands started 7 million years ago and thus coincides with the emergence of a large proportion of insular shrubby plants (80% of insular lineages on the Canary Islands are woody^40,41^). The higher stem specific density of shrubs compared to herbaceous species decreases the risk of hydraulic failure^41,42^. Succulence represents another key adaptation to the arid environments and this trait is also well represented on the Canary Islands^43^, and this is here captured by leaf thickness. Including Leaf thickness into the trait space (Fig. 2a) highlights the importance of succulence in response to aridity.

The dominance of shrubby species on Tenerife is consistent with the idea that small herbaceous colonisers gradually evolve into taller plants with increased stem density to avoid competition with other species^44^. Yet, the evolution of several Tenerife shrub species and the subsequent increase in shrubbiness has mostly occurred in steep canyons and high-elevation ecosystems^45^, where conditions are unfavourable for trees and competition with taller plants is low. The underrepresentation of tall plants in Tenerife’s trait space reflects the comparatively low number of tree species (mainly in laurel and pine forest, and thermophilous woodland ecosystems) compared to shrubs. Overall, the functional dominance of shrubs in the flora of Tenerife is evidence of shrubbiness as a general island syndrome.

### Biogeography and evolution have jointly shaped island plant trait space

We analysed the relationship of eight plant functional traits (Fig. 2a and Extended Data Fig. 2) for the five species groups (Fig. 2b-f) and found that trait spaces of almost all groups were highly aggregated in the centre of the island trait space. The trait space of non-endemic natives (Fig. 2b), encompasses a large range of trait combinations, which relative to other groups, are evenly distributed across the island trait space, i.e., from light-seeded, short plants to tall plants with heavy seeds (ranging from 0.01 - 230 mg seed mass and 0.1 - 20 m plant height). Canary endemic species (Fig. 2d) also have a large range of trait combinations, but the trait space is highly dominated by shrubs and extends towards species with high leaf thickness, a trait associated with drought tolerance, and towards species with nitrogen rich leaves. The trait space of Canary endemics also extends towards species with contrasting trait combinations, i.e., low leaf thickness and low leaf nitrogen content, that are consistent with a conservative life strategy. The trait space of Tenerife endemics (Fig. 2e) is mainly dominated by shrubs and succulent species. In contrast, the trait space of Macaronesian endemics (Fig. 2c) has a bimodal distribution associated with the prevalence of both shrubs and trees. Trait combinations of tree species within the Macaronesian endemics, e.g., *Laurus novocanariensis* with its large stature and seed mass (25 m and 600 mg, respectively), have not emerged from *in-situ* speciation but rather from the relictualization of laurel forests, an ecosystem that is largely extinct on the mainland^38^. Lastly, cladogenetic species (Fig. 2f) occur also in the centre of the island trait space, with most species having intermediate stature, light seeds, and thick leaves with low leaf dry matter content (typical for succulents). Trait spaces of cladogenetic species and Tenerife endemics are similar, as most Tenerife endemics emerged via cladogenesis.

### Biogeography and evolution leave imprints on island functional diversity

To capture different aspects of trait distributions among all five groups, we calculated three components of functional diversity: functional richness, functional evenness and functional dispersion^23^ (Fig. 3). Further, we assessed the contribution, i.e., whether a group increases island trait space or not, and the uniqueness of each group within the island trait space by calculating functional contribution and originality^23^ (Extended Data Fig. 4).

**Figure 3.**
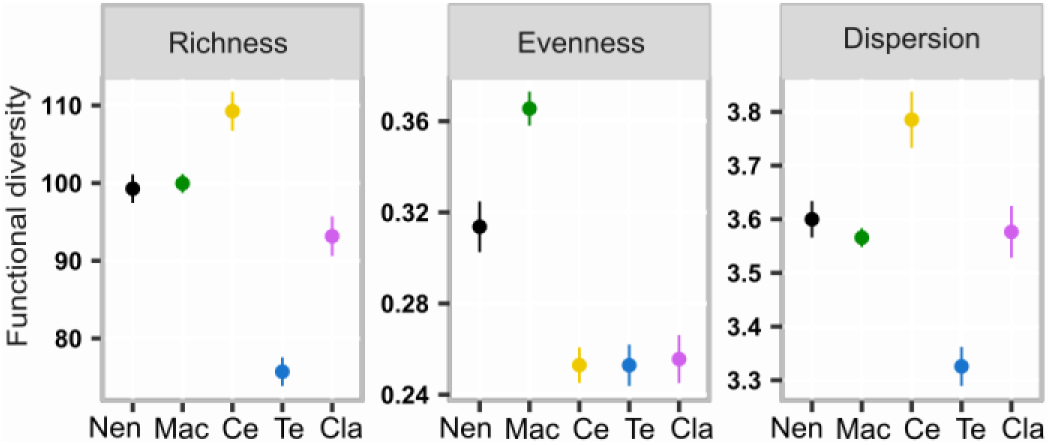
Contribution of biogeographical and evolutionary processes to the functional diversity of Tenerife. Functional richness, functional dispersion and functional evenness were calculated using n-dimensional hypervolumes while controlling for the number of species richness per group (see Methods). Different colours refer to a species group, i.e., black to non-endemic native (Nen), green to Macaronesian endemics (Mac), yellow to Canary endemics (Ce), blue to Tenerife endemics (Te) and purple to cladogenetic species (Cla). Dots and error bars correspond to the mean values and 95% confidence intervals, based on species-richness based rarefaction values.

We found that functional richness (Fig. 3), i.e., the total amount of trait space^20^, and functional dispersion (Fig. 3), i.e., the functional divergence or convergence of species within a trait space, exhibit a similar pattern among groups and are highest for Canary endemics. This result, together with the significant functional contribution and originality of the Canary endemics (Extended Data Fig. 4), indicates that this group possesses unique trait combinations and causes an expansion of the trait space of Tenerife. In contrast, we found that Tenerife endemics have the lowest functional richness and dispersion among all groups (Fig. 3). This contrasting pattern emerges because several Canary endemics occupy the margins of the trait space (i.e., small herbs, tall shrubs and trees), whereas Tenerife endemics are mainly found in the centre of island trait space (i.e., shrubs and small succulents) (Fig. 2 and Fig. 3). Canary endemic species at the margins of Tenerife’s trait space, such as *Spartocytisus supranubius*, a 4-metre-tall shrub with dense stems (stem specific density = 0.7 mg mm^−3^) or *Monanthes laxiflora*, a small succulent (0.2 m tall and 0.4 mm thick leaves), are examples of species with unique trait combinations that expand the island trait space. The high functional diversity of Canary endemics is consistent with the idea that insular species evolve new morphological characteristics to take advantage of unoccupied niche space, thereby avoiding interspecific competition^46^. The high habitat diversity of the Canary Islands provides ample opportunities for species that, together with inter-island dispersal, promotes speciation and drives the high functional diversity of the Canary endemics. Conversely, Tenerife endemics are functionally quite similar; these species occupy similar island habitats, have a low degree of niche differentiation, and have likely emerged via allopatric speciation^45^.

Functional evenness (Fig. 3), i.e., the regularity of species’ distribution across trait space was highest for Macaronesian endemics and non-endemic natives. The high evenness values for these groups indicate that species are equally distributed among trees, shrubs, and herbs. Further, the high functional contribution and functional originality of Macaronesian endemics (Extended Data Fig. 4) illustrates that certain species and relictualization events may contribute disproportionately to island trait space by adding unique trait combinations.

Unexpectedly, we found that cladogenetic species did not expand the trait space of Tenerife (cf. intermediate functional richness and dispersion values in Fig. 3). We attribute this result to a greater frequency of non-adaptive speciation events relative to adaptive ones^47^. Unlike adaptive radiations, where species commonly adapt trait values to cope with changing environmental conditions^32^, non-adaptive radiations^47^ may not result in shifts in trait values because environmental conditions may be similar between isolated populations. This leads to newly evolved species that are functionally similar to ancestral species. Further, the uneven trait space of cladogenetic species (cf. low functional evenness values in Fig. 3) is consistent with the idea that environmental filters result in convergence around similar trait combinations^48^. The aridification of the Canary Islands created new habitats, yet the harsh environmental conditions of these habitats limit viable trait combinations^49^. Lastly, non-endemic native species had high functional richness and dispersion values across groups (Fig. 3). Contrary to our expectation, this result suggests that long-distance dispersal is contributing to functional diversity rather than constraining it.

### How radiated lineages contribute to the island functional trait space

To understand how speciation due to radiations has shaped the functional trait space of Tenerife, we quantified the functional contribution and originality of the 21 major lineages (i.e., 161 species, cf. Extended Data Fig. 5-6), which have radiated in Macaronesia and are present on the Canary Islands and Tenerife, to the island trait space. We found that the most diverse radiated lineages, *Aeonium* (>30 species) and *Polycarpaea* lineage (7 species), contributed significantly to the expansion of Tenerife trait space. Both lineages expand the trait space towards small plants with thick leaves (Extended Data Fig. 5c). In contrast, the vast majority of radiated lineages did not increase island trait space. This result suggests that species within the *Aeonium* and *Polycarpaea* lineages evolved trait combinations including succulent leaves and remarkable size changes that allow them to exploit available niche space across a variety of habitats. The minimal contributions of the other, less diverse radiated lineages to island trait space might indicate that these radiations are to a large degree non-adaptive. Yet, it could also indicate that while the newly evolved species are similar to those of co-occurring species, they may have evolved other traits not measured in this study that enable them to exploit empty niche space.

We conclude that evolution via different speciation pathways (i.e., anagenesis and cladogenesis) at the archipelago level, together with biogeographical processes such as relictualization and inter-island dispersal at the archipelago or meta-archipelago level, have jointly shaped the trait space of Tenerife. Contrary to our hypothesis, cladogenesis has led to functional convergence, and therefore only marginally to increases functional diversity. The predominance of shrubbiness and leaf succulence across the island flora additionally reflects a strong imprint of environmental filtering acting on the trait space of Tenerife. Overall, our results show how functional diversity of an oceanic island flora emerges from the interaction among biogeographical, ecological and evolutionary processes. Our approach offers a first step towards understanding, from a trait-based perspective, the assembly of an entire native flora.

## Methods

### Tenerife as a model system

Tenerife is the largest (2058 km^2^) and tallest (3715 m a.s.l.) of the Canary Islands, an archipelago that belongs to the meta-archipelago Macaronesia, a floristic region off the coast of Northern Africa and Western Europe in the Atlantic Ocean^33^. The island is of volcanic origin and its oldest substrate is about 8 million years old^50^. Due to its complex topography, dynamic geological history^50^, and interaction with the north-eastern trade wind system, Tenerife has a very high environmental heterogeneity and a wide range of different ecozones and habitats^51^.

### Floristic and biogeographical data

We analysed all native seed plant species of Tenerife as listed in the latest version of the vascular plant species checklist of the Canary Islands^35^. Since the native status of several Canary Islands species remains unresolved^36^, we included only species categorised as native (i.e., “*nativa segura*” status from the Canary Islands plant checklist), yielding a total of 436 species^35^. We classified species into five different groups according to species endemism status from the checklist^35^ as follows: non-endemic native (78 species), Macaronesian endemics (45 species), Canary Islands endemics (186 species) and endemics to Tenerife (127 species). The cladogenetic species group (264 species) is composed of 21 lineages that have radiated across the Canary Islands and Macaronesia and have produced more than three extant species (according to^36^); therefore, this group contains Macaronesian, Canary as well as Tenerife endemics.

### Trait sampling and measurement

We collected and measured leaf and stems traits^16,18^ for 79% and 76 % respectively, of all Tenerife native seed plants (344 and 330 species): leaf area (mm2) as the one-side surface area of the individual lamina, leaf dry matter content (LDMC; mg g^−1^) as the leaf dry mass per unit of water-saturated fresh mass, leaf mass per area (LMA; g m^−2^) as the leaf dry mass per unit of lamina surface area, leaf Nitrogen content (Leaf N_s_; mg g^−1^) as the nitrogen content per unit of lamina dry mass, leaf thickness (Leaf th; mm), stem specific density (Stem density; mg mm^−3^) as dry mass per unit of fresh stem volume. We sampled plants across the entire island at more than 500 locations (Extended Data Fig. 7), covering the full range of elevational and climatic gradients of Tenerife from sea level to 2700 m a.s.l. We sampled rare species (when possible) at only one site for conservation reasons and a few rare species (n = 20) in botanical gardens^52^ (Jardín de Aclimatación de La Orotava and Jardines Campus Universidad de La Laguna in Tenerife, and Jardín Botánico Canario Viera y Clavijo in Gran Canaria). We confirmed species identity with botanical experts, Rüdiger Otto and Rubén Barone, and identification books^53^. We collected three replicates (individuals per species) for 60% of all 361 sampled species, one to two replicates for 16%, and four to five replicates for 6% of all the sampled species.

Trait values for leaves, stems and seeds were measured following standardised protocols^54^. We collected healthy, fully expanded sun leaves from individual plants. Depending on leaf size, we collected between 10 (for > 1 cm^2^ leaves) to 100 (for < 1 cm^2^ leaves) leaves per individual. To measure stem traits and ensure that plants were not damaged, we collected samples from the first adjacent branch of the main plant stem, when plants were shrubs. For small herbs and vines, we collected the plant organ acting as a stem. To sample the stem density of trees, we used an increment borer, to extract a wood core from the bark inward (10 mm diameter core taken at ~1.2 m above ground). We stored the fresh plant material in coolers to prevent dehydration and measured fresh leaf mass using an analytical balance (0.01 mg precision from PCB 2500-2 Kern & Sohn) within 24 hours after collection. Leaf thickness and leaf area (leaves smaller than 1 cm^2^ were scanned at 600 dpi and leaves larger than 1 cm^2^ at 300 dpi) were also measured within 24 hours after collection. Leaf area was calculated using WinFOLIA software (Version 2016b Pro, Regent Instruments Canada, 2016). To measure the volume of fresh stems, we first measured their length and diameter with a digital calliper. As stems are not perfect cylinders, we measured diameter in three different stem sections and used the mean value per stem. We computed fresh stem volume using the following formula for cylinders: V=Πr^2^h, where Π is the constant Pi, r is radio, and h is height. We oven dried leaves and stems for 48 hours, or until a stable weight was reached, at 80°C and then measured leaf and stem dry mass using the same analytical balance. Nitrogen content of the dry leaves was determined by a C/N elemental analyser (Vario EL III, elementar, Hanau, Germany).

We measured seed mass (mg) for 74% (322 species) of all native seed plants of Tenerife as the dried mass of an individual seed in the field (9 species) and at the seed bank from the Jardín Botánico Canario “Viera y Clavijo” in Gran Canaria (313 species). We counted between 5 to 200 seeds per species and weighed them using an analytical balance (0.001 mg precision). We obtained individual seed mass by dividing the total mass of the seeds by the number of seeds. For very small seeds (< 0.1 mm), we calculated seed mass using a test tube containing a volume of seeds for which the seed count was known. We obtained maximum plant height (Height; m), which is the upper boundary of the main photosynthetic tissue at maturity in metres, for 97% of all native species of Tenerife from the literature^53^.

Prior to analysis, we inspected the density distribution of single traits and correlations among traits (Extended Data Fig. 3). We then used phylogenetic trait imputation to estimate missing trait values. We imputed trait values for 20% of leaf trait values, 27% of stem density, 26% of seed mass, and 3% of maximum plant height. We followed the phylogenetic imputation procedure suggested by^55^. To this end, we first constructed the phylogeny using the mega-phylogeny^56^ and conservatively bound species to the backbone using dating information from congeners in the tree with the ‘congeneric.merge’ function in package ‘pez’^57^ implemented for R software^58^. After checking for synonyms, we bound 430 species to the phylogeny. We used a random forest algorithm using the ‘missForest’ function from the MissForest R package^59^ to predict missing trait values for the 430 species (see density distribution of original and imputed trait values in Extended Data Fig. 3b). We filled in the missing six species trait values using the results of a naive prediction without phylogenetic information, and which used the random forest algorithm. Using out-of-bag error, we measured the prediction error of the random forest algorithm that included phylogenetic relationships among species against naive predictions. We found that phylogenetically informed imputation performed better than the naive imputation for nearly all traits (Extended Data Table 1).

### Global trait data

For comparison of the Tenerife trait space and the global traits space, we used the global trait data from^18^, which has complete information for six plant traits (Leaf area, LMA, Leaf N, Seed mass, Stem density and Height) for 2213 plant species. Before comparison among Tenerife and global trait space, we removed 14 species from the global trait data that belonged to the flora of Tenerife. We compared 2199 species from the global trait data with 436 Tenerife species.

### Functional trait space

We performed a principal component analysis (PCA) on the log- and z-transformed (centred and rescaled to unit variance) mean trait values. For the comparison of Tenerife versus global trait space of plant form and function (Fig. 2), we used only six plant functional traits: leaf area, LMA, leaf N, seed mass, stem density and height. For the trait space analysis of Tenerife island and the different biogeographical groups (Fig. 3a and Extended Data Fig. 2), we used eight plant functional traits: leaf area, LMA, LDMC, leaf N, leaf th, seed mass, stem density and height. We visualised the trait space of each group based on the eight plant traits (Fig. 3).

### Functional diversity

We calculated three components of functional diversity (Fig. 3) using the n-dimensional functional hypervolumes by^60^. This approach to functional diversity is thought to be more accurate than traditional approaches because it accounts for gaps in trait space and, in doing so, avoids overestimation of functional diversity. To compute hypervolumes, we used a fixed kernel bandwidth for all groups using the ‘estimate_bandwidth’ (using cross-validation as kernel estimator method) function in the R package Hypervolume^61^. We used the Gaussian method to build hypervolumes, as it is the least sensitive method to variation in bandwidth and fits the data loosely, which is suitable for functional diversity and fundamental niche modelling applications^60^. Using the hypervolumes, we then calculated functional richness, evenness and dispersion using the function ‘kernel.alpha’, ‘kernel.evenness’, and ‘kernel.dispersion’ in the R package BAT^23^. Functional richness is the total volume of a trait space. Functional dispersion quantifies how spread or dense a given trait space is, by calculating the average difference between the trait space centroid and random points (i.e., randomly placed species) within the boundaries of the hypervolume^23^. Functional evenness quantifies how regular a given trait space is, by calculating the overlap between the observed hypervolume and a theoretical, perfectly even hypervolume^23^. Sørensen (volume of the intersection of island trait space and global trait space, divided by the volume of union of island trait space and global trait space) similarity coefficients (cf. Fig. 2), by first building Gaussian hypervolumes with a fixed bandwidth, for both global and island data and then computed hypervolume overlap statistics, using the ‘hypervolume_overlap_statistics’ function from the R package Hypervolume^60^. To estimate the contribution (whether a group increases island trait space or not) and originality (a group with high originality value has a unique within the island trait space, which translates into unique trait combinations relative to the island) of the five groups to the island trait space, we first calculated functional contribution^23^ as the net contribution of each single species to the total island hypervolume, and functional originality^23^, as the average dissimilarity difference between a given species and a sample of random points (10% of the total random points) within the boundaries of the island hypervolume^23^. To assess the functional contribution and originality of each group, we plotted the functional contribution and functional originality values per of the species belonging to a given group against all other species that do not belong to the group as box plots (Extended Data Fig. 4).

### Data analysis

We used Kruskal-Wallis tests to assess differences among the first and second principal components of the trait spaces of each group (Extended Data Fig. 2c), as well as for assessing the statistical significance of functional contribution and originality of each group (Extended Data Fig. 4) and 21 radiated lineages (Extended Data Fig. 5 and Extended data Table 2) to the island trait space. For the test, we used the ‘kruskal’ function in the R package Agricolae^62^. Because functional diversity metrics are commonly affected by the number of species^63^, we performed species-richness based rarefaction to ensure that values were comparable across the five groups. To this end, we selected a minimum common number of species (n= 30) across the five groups, which we randomly sampled 99 times with replacement, and calculated the functional diversity metrics each time per group. We computed mean values and 95% confidence intervals from all samples to compare functional diversity metrics across groups (Fig. 3). Lastly, we visualised the functional trait spaces (Fig. 2) using the ‘stat_density_2d’ function in the R package ggplot2^64^. We performed all statistical analysis and data visualisation using R software version 4.1.0.

### Data Availability

The trait data and floristic information of the species that support the findings of this study are available in Figshare repository https://figshare.com/s/a9d343529990afa0c799

### Code Availability

The analysis and data visualisation performed in R software that support the findings of this study are available in Figshare repository https://figshare.com/s/a9d343529990afa0c799

## Acknowledgements

M.P.B.B. and H.K. acknowledge funding from the German Research Foundation (DFG) Research Training Group 1644 ‘Scaling Problems in Statistics’, grant no. 152112243. D.C. acknowledges funding from the Agencia Nacional de Investigación y Desarrollo (Chile; FONDECYT Regular No 1201347). We thank the Jardín Botánico Canario “Viera y Clavijo” in Gran Canaria for allowing measurements in the seed bank as well as for the samples taken of rare species that were not possible to find in the field. We thank Nora Strassburger, Mercedes Vidal Rodríguez and Arnau Andreu Diez for invaluable assistance in the field and in the laboratory. We also thank Rubén Barone for assisting with plant identification.

## Author contributions

H.K. conceived the initial idea. M.P.B.B., D.C., P.W., and H.K. further developed the concepts and designed the research. M.P.B.B. collected and measured plant trait data and performed the analysis. P.D. supported the analysis. R.O. supported plant trait data collection and species identification. All authors contributed to the interpretation of the results and writing of the paper.

## References

1. Carlquist, S. The biota of long-distance dispersal. II. Loss of dispersibility in Pacific Compositae. Evolution. 20, 30–48 (1966).

2. Darwin, C. On the Origin of Species. (Murray, London, 1859).

3. Burns, K. C. Evolution in isolation: the search for an island syndrome in plants. (Cambridge University Press, 2019).

4. Patiño, J. et al. A roadmap for island biology: 50 fundamental questions after 50 years of The Theory of Island Biogeography. J. Biogeogr. 44, 963–983 (2017).

5. Whittaker, R. J., Fernández-Palacios, J. M., Matthews, T. J., Borregaard, M. K. & Triantis, K. A. Island biogeography: Taking the long view of nature’s laboratories. Science 357, eaam8326 (2017).

6. Losos, J. B. & Ricklefs, R. E. Adaptation and diversification on islands. Nature 457, 830–836 (2009).

7. Craven, D., Knight, T. M., Barton, K. E., Bialic-Murphy, L. & Chase, J. M. Dissecting macroecological and macroevolutionary patterns of forest biodiversity across the Hawaiian archipelago. Proc. Natl. Acad. Sci. 116, 16436–16441 (2019).

8. MacArthur, R. H. & Wilson, E. O. The Theory of Island Biogeography. The Journal of Wildlife Management vol. 33 (Princeton, NJ: Princeton University Press, 1969).

9. Gillespie, R. G. & Baldwin, B. G. ‘Island biogeography of remote archipelagoes.’ The theory of island biogeography revisited. Princeton University Press (2010).

10. Weigelt, P. et al. Global patterns and drivers of phylogenetic structure in island floras. Sci. Rep. 5, 1–13 (2015).

11. Loiseau, N. et al. Global distribution and conservation status of ecologically rare mammal and bird species. Nat. Commun. 11, 5071 (2020).

12. Schrader, J., Wright, I. J., Kreft, H. & Westoby, M. A roadmap to plant functional island biogeography. Biol. Rev. 96, 2851–2870 (2021).

13. Violle, C., Reich, P. B., Pacala, S. W., Enquist, B. J. & Kattge, J. The emergence and promise of functional biogeography. Proc. Natl. Acad. Sci. U. S. A. 111, 13690–13696 (2014).

14. Lavorel, S. & Garnier, E. Predicting changes in community composition and ecosystem functioning from plant traits: Revisiting the Holy Grail. Funct. Ecol. 16, 545–556 (2002).

15. Violle, C. et al. Let the concept of trait be functional! Oikos 116, 882–892 (2007).

16. Wright, I. J. et al. The worldwide leaf economics spectrum. Nature (2004) doi:10.1038/nature02403.

17. Reich, P. B. The world-wide ‘fast-slow’ plant economics spectrum: A traits manifesto. J. Ecol. 102, 275–301 (2014).

18. Díaz, S. et al. The global spectrum of plant form and function. Nature 529, 167–171 (2016).

19. Díaz, S. & Cabido, M. Vive la diffèrence: Plant functional diversity matters to ecosystem processes. Trends Ecol. Evol. 16, 646–655 (2001).

20. Mouchet, M. A., Villéger, S., Mason, N. W. H. & Mouillot, D. Functional diversity measures: An overview of their redundancy and their ability to discriminate community assembly rules. Funct. Ecol. 24, 867–876 (2010).

21. Wieczynski, D. J. et al. Climate shapes and shifts functional biodiversity in forests worldwide. Proc. Natl. Acad. Sci. 116, 7591–7591 (2019).

22. Bruelheide, H. et al. Global trait–environment relationships of plant communities. Nat. Ecol. Evol. 2, 1906–1917 (2018).

23. Mammola, S. & Cardoso, P. Functional diversity metrics using kernel density n-dimensional hypervolumes. Methods Ecol. Evol. 11, 986–995 (2020).

24. König, C. et al. Biodiversity data integration—the significance of data resolution and domain. PLoS Biol. 17, 1–16 (2019).

25. Cornwell, W. K., Pearse, W. D., Dalrymple, R. L. & Zanne, A. E. What we (don’t) know about global plant diversity. Ecography (Cop.). 42, 1819–1831 (2019).

26. Heleno, R. H. & Vargas, P. How do islands become green? Glob. Ecol. Biogeogr. 24, 518–526 (2015).

27. Barajas-Barbosa, M. P., Weigelt, P., Borregaard, M. K., Keppel, G. & Kreft, H. Environmental heterogeneity dynamics drive plant diversity on oceanic islands. J. Biogeogr. 47, 2248–2260 (2020).

28. Stuessy, T. & Crawford, D. J. Patterns of Phylogeny in the Endemic Vascular Flora of the Juan Fernandez Islands, Chile. Am. Soc. Plant Taxon. 15, 338–346 (1990).

29. Stuessy, T. F. et al. Anagenetic evolution in island plants. J. Biogeogr. 33, 1259–1265 (2006).

30. Emerson, B. C. & Gillespie, R. G. Phylogenetic analysis of community assembly and structure over space and time. Trends Ecol. Evol. 23, 619–630 (2008).

31. Bohle, U. R., Hilger, H. H. & Martin, W. F. Island colonization and evolution of the insular woody habit in Echium L. (Boraginaceae). Proc. Natl. Acad. Sci. 93, 11740–11745 (1996).

32. Givnish, T. J. et al. Origin, adaptive radiation and diversification of the Hawaiian lobeliads (Asterales: Campanulaceae). Proc. R. Soc. B Biol. Sci. 276, 407–416 (2009).

33. Florencio, M. et al. Macaronesia as a Fruitful Arena for Ecology, Evolution, and Conservation Biology. Front. Ecol. Evol. 9, (2021).

34. Fernández-Palacios, J. M. Climatic responses of plant species on Tenerife, The Canary Islands. J. Veg. Sci. 3, 595–603 (1992).

35. Acebes-Ginovés, J., León Arencibia, M. & Rodríguez Navarro, M. Lista de especies silvestres de Canarias. Hongos, plantas y animales terrestres 2009. (Santa Cruz de Tenerife: Gobierno de Canarias, 2010).

36. Price, J. P. et al. Colonization and diversification shape species–area relationships in three Macaronesian archipelagos. J. Biogeogr. 45, 2027–2039 (2018).

37. De Nascimento, L., Willis, K. J., Fernández-Palacios, J. M., Criado, C. & Whittaker, R. J. The long-term ecology of the lost forests of la Laguna, Tenerife (Canary Islands). J. Biogeogr. 36, 499–514 (2009).

38. Fernández-Palacios, J. M. et al. A reconstruction of Palaeo-Macaronesia, with particular reference to the long-term biogeography of the Atlantic island laurel forests. J. Biogeogr. 38, 226–246 (2011).

39. Carlquist, S. J. Island biology. (Columbia University Press, 1974).

40. van Huysduynen, A. et al. Temporal and palaeoclimatic context of the evolution of insular woodiness in the Canary Islands. Ecol. Evol. 11, 12220–12231 (2021).

41. Dória, L. C. et al. Insular woody daisies ( Argyranthemum, Asteraceae) are more resistant to drought-induced hydraulic failure than their herbaceous relatives. Funct. Ecol. 32, 1467–1478 (2018).

42. Lens, F., Davin, N., Smets, E. & del Arco, M. Insular Woodiness on the Canary Islands: A Remarkable Case of Convergent Evolution. Int. J. Plant Sci. 174, 992–1013 (2013).

43. Shmida, A. & Werger, M. J. A. Growth form diversity on the Canary Islands. Vegetatio 102, 183–199 (1992).

44. Biddick, M., Hendriks, A. & Burns, K. C. Plants obey (and disobey) the island rule. Proc. Natl. Acad. Sci. 116, 17632–17634 (2019).

45. Fernández-Palacios, J. M. et al. Evolutionary winners are ecological losers among oceanic island plants. J. Biogeogr. 48, 2186–2198 (2021).

46. Silvertown, J. The Ghost of Competition Past in the Phylogeny of Island Endemic Plants. 92, 168–173 (2004).

47. Rundell, R. J. & Price, T. D. Adaptive radiation, nonadaptive radiation, ecological speciation and nonecological speciation. Trends Ecol. Evol. 24, 394–399 (2009).

48. Spasojevic, M. J. & Suding, K. N. Inferring community assembly mechanisms from functional diversity patterns: The importance of multiple assembly processes. J. Ecol. 100, 652–661 (2012).

49. Carvajal-Endara, S., Hendry, A. P., Emery, N. C. & Davies, T. J. Habitat filtering not dispersal limitation shapes oceanic island floras: species assembly of the Galápagos archipelago. Ecol. Lett. 20, 495–504 (2017).

## Methods references

50. Troll, V. R. & Carracedo, J. C. The Geology of Tenerife. in The Geology of the Canary Islands 227–355 (Elsevier, 2016).

51. Fernández-Palacios, J. M. & Nicolás, J. P. Altitudinal pattern of vegetation variation on Tenerife. J. Veg. Sci. (1995).

52. Perez, T. M. et al. Botanic gardens are an untapped resource for studying the functional ecology of tropical plants. Philos. Trans. R. Soc. B Biol. Sci. 374, 20170390 (2019).

53. Muer, T., Sauerbier, H. & Calixto, F. C. Die Farn-und Blütenpflanzen der Kanarischen Inseln. (Margraf Publishers GmbH, 2016).

54. Pérez-Harguindeguy, N. et al. Corrigendum to: New handbook for standardised measurement of plant functional traits worldwide. Aust. J. Bot. 64, 715 (2016).

55. Penone, C. et al. Imputation of missing data in life-history trait datasets: which approach performs the best? Methods Ecol. Evol. 5, 961–970 (2014).

56. Smith, S. A. & Brown, J. W. Constructing a broadly inclusive seed plant phylogeny. Am. J. Bot. 105, 302–314 (2018).

57. Pearse, W. D. et al. An introduction to pez. https://cran.r-project.org/web/packages/pez/ (2021).

58. R Core Team. R: A language and environment for statistical computing. R Foundation for Statistical Computing, Vienna, Austria. (2021).

59. Stekhoven, D. J. & Bühlmann, P. missForest: Nonparametric Missing Value Imputation using Random Forest R package version 1.3. https://cran.r-project.org/web/packages/missForest/index.html (2013).

60. Blonder, B. Hypervolume concepts in niche- and trait-based ecology. Ecography. 41, 1441–1455 (2018).

61. Blonder, B. R Pakcage: Hypervolume: High Dimensional Geometry and Set Operations Using Kernel Density Estimation, Support Vector Machines, and Convex Hulls. https://cran.r-project.org/web/packages/hypervolume/ (2021).

62. De Mendiburu, F. Agricolae: statistical procedures for agricultural research. R package version, 1(1), 1–4. https://cran.r-project.org/web/packages/agricolae/ (2021).

63. Schleuter, D., Daufresne, M., Massol, F. & Argillier, C. A user’s guide to functional diversity indices. Ecol. Monogr. 80, 469–484 (2010).

64. Wickham, H. ggplot2: elegant graphics for data analysis. Springer-Verlag (2016).

