## Supplemental Material for "Assembly of functional diversity in an oceanic island flora"

### Extended data figures and tables

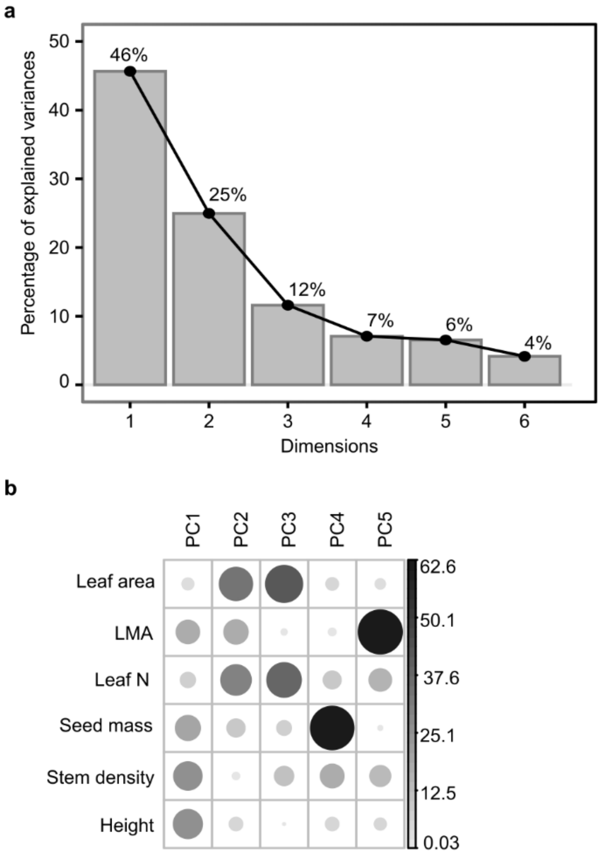


**Extended data Fig. 1 | Overview of the principal components and explained variance of both island and global data a.** Percentage of explained variance of the six dimensions of the principal component analysis (PCA) based on six plant functional traits of Tenerife's native seed plants and seed plants from across the globe (2199 species). **b.** Percentage of contributions of each trait to each dimension of the PCA. Trait values were z- and log-transformed prior to the PCA.

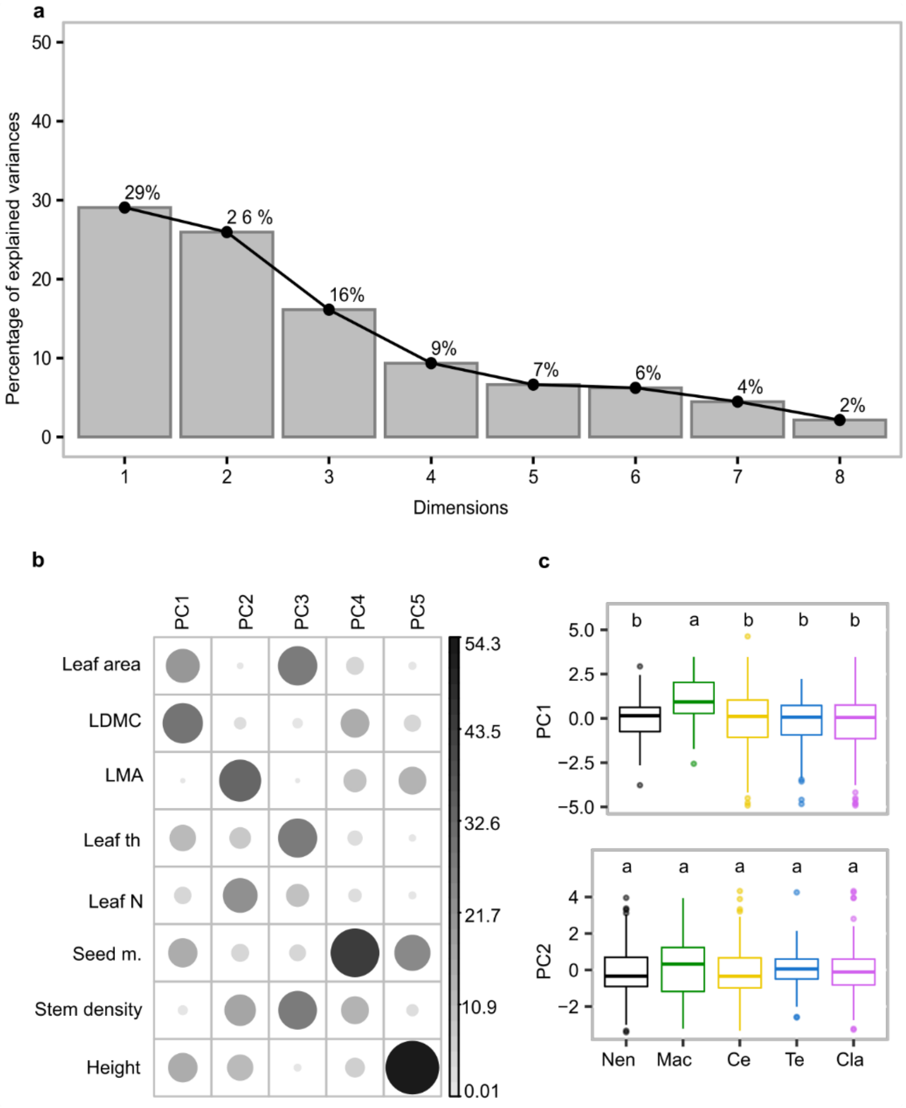


**Extended data Fig. 2 | Overview of principal components analysis (PCA) for eight plant functional traits (436 species) of Tenerife native seed plants**. **a.** Percentage of explained variance of the eight dimensions of the PCA. **b.** Percentage of contributions of each trait to each dimension of the PCA. Trait values were log- and z-transformed. **c.** Differences among the first and second principal component of the trait spaces of different biogeographical groups, Non-endemic natives (Nen), Macaronesian endemics (Mac), Canary Islands endemics (Ce), Tenerife endemics (Te) and cladogenetic species (Cla). Identical letters indicate no significant differences (Kruskal-Wallis text, P>0.05).

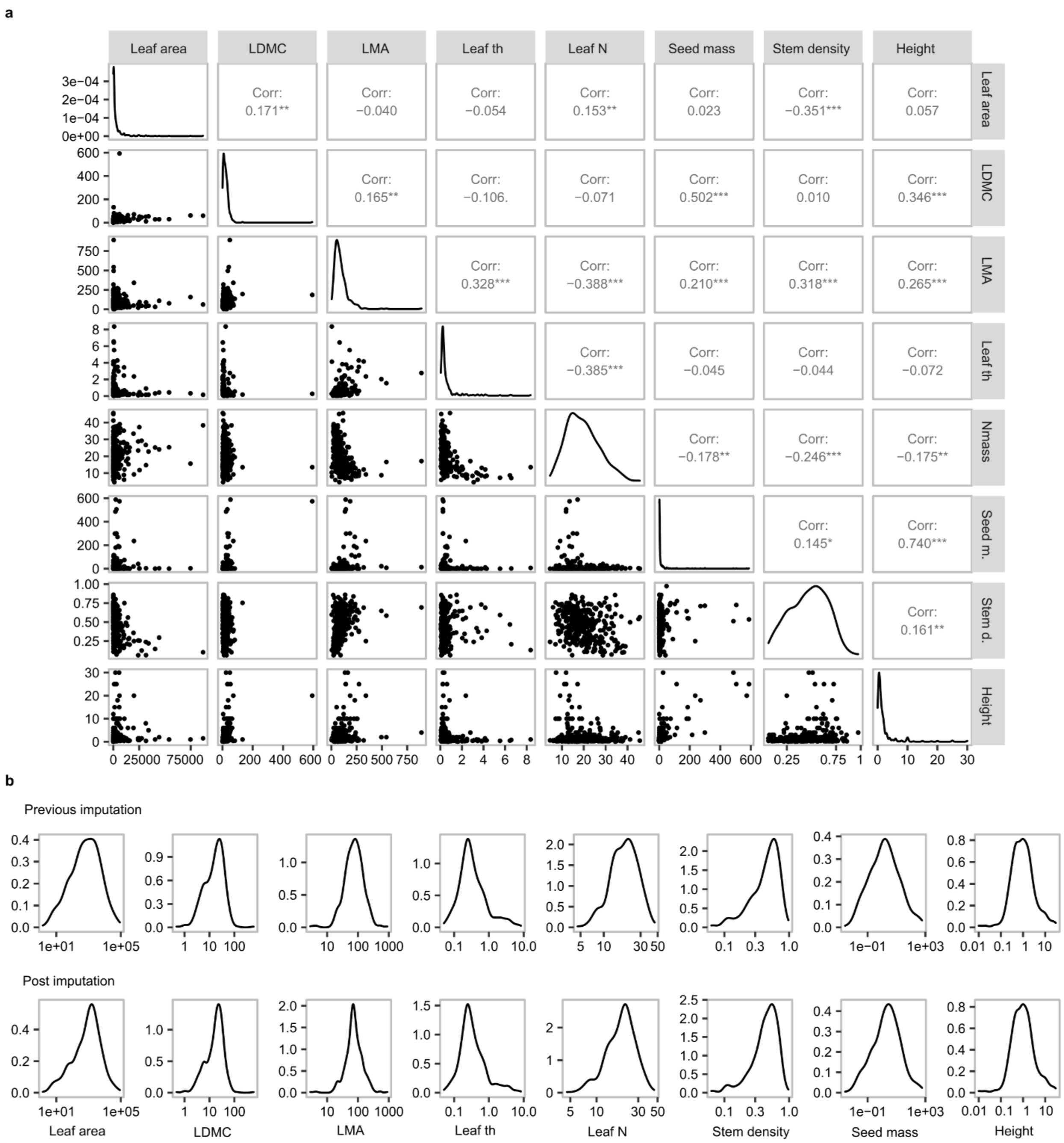


**Extended data Fig. 3 | Trait data overview and assessment. a.** Pearson correlation coefficients among eight plant traits and their density distribution, which we collected for approximately 80 % of all native seed plant species of Tenerife. **b.** Density distribution of the original trait values only, and including imputed trait values for each trait. Percentages correspond to the proportion of species with imputed trait values (n = 436 native plant species).

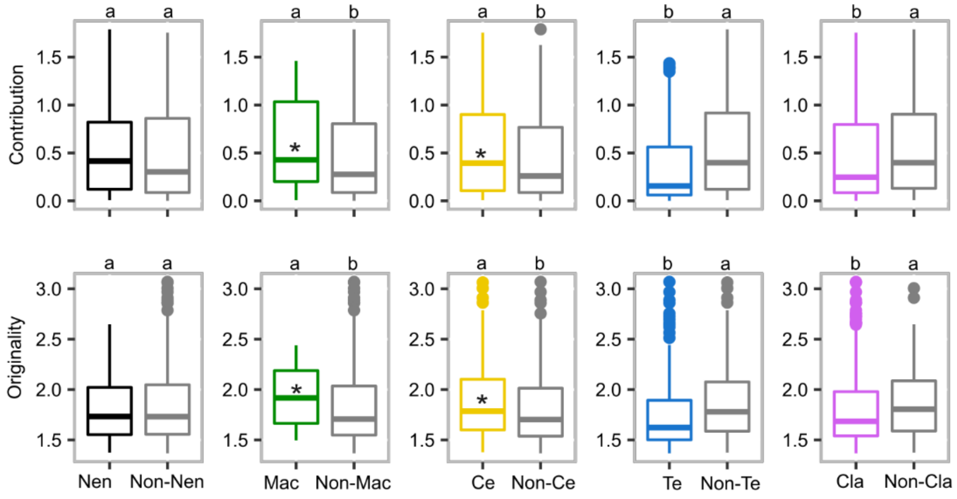


**Extended data Fig. 4 | Functional contribution and originality for the five biogeographical groups**. Each colour refers to a group, black to Non-endemic native (Nen), green to Macaronesian endemics (Mac), yellow to Canary endemics (Ce), blue to Tenerife endemics (Te) and purple to cladogenetic species (Cla). “Non” preceding a group abbreviation, e.g., “Non-Nen” indicates all island species except for non-endemic native species. Dots and error bars correspond to the mean values and 95% confidence intervals. Asterisks (*) indicate that biogeographical groups have significantly higher functional contribution or functional originality (where, identical letters indicate no significant differences Kruskal-Wallis text, P>0.05). Functional contribution and originality are based on hypervolume calculations, which were estimated using eight plant functional traits sampled for 436 native seed plant species of Tenerife.

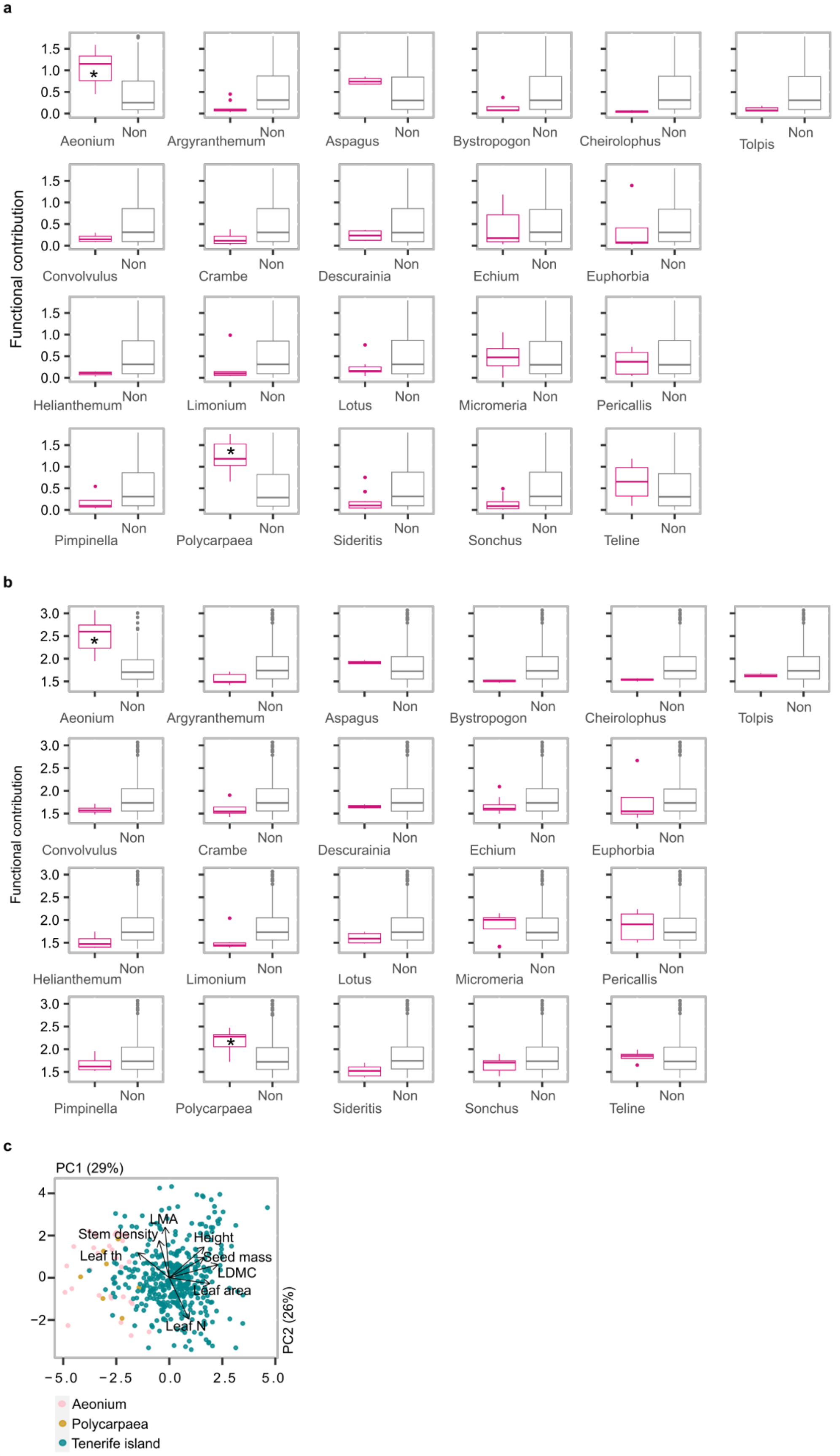
a
Functional contribution
1.5
1.0
0.5
0.0
Aeonium
Argyranthemum
Aspagus
Bystropogon
Cheirolophus
Tolpis
Convolvulus
Crambe
Descurainia
Echium
Euphorbia
Helianthemum
Limonium
Lotus
Micromeria
Pericallis
Pimpinella
Polycarpaea
Sideritis
Sonchus
Teline
Non
*
b
Functional contribution
3.0
2.5
2.0
1.5
Aeonium
Argyranthemum
Aspagus
Bystropogon
Cheirolophus
Tolpis
Convolvulus
Crambe
Descurainia
Echium
Euphorbia
Helianthemum
Limonium
Lotus
Micromeria
Pericallis
Pimpinella
Polycarpaea
Sideritis
Sonchus
Teline
Non
*
c
PC1 (29%)
PC2 (26%)
LMA
Stem density
Height
Leaf th
Seed mass
LDMC
Leaf area
Leaf N
4
2
0
-2
-5.0
-2.5
0.0
2.5
5.0
Aeonium
Polycarpaea
Tenerife island

**Extended data Fig. 5 | Functional contribution and originality of the major radiated plant lineages present in Tenerife**. **a**. Functional contribution and **b.** functional originality for the 21 major radiated lineages. Note that only *Aeonium* alliance and *Polycarpaea* lineage (marked with *) are significantly contributing to the expansion of Tenerife trait space, as their functional contribution and originality are significantly different from other species that do not belong to the lineage (results based on Kruskal-Wallis, with alpha level 0.05). “Non” indicates all island seed plant native species, except for species included in the lineage displayed in a single boxplot. **c.** Location of *Aeonium* and *Polycarpaea* species in the island trait space; both lineages extend it towards high values of leaf thickness (i.e., Leaf th).

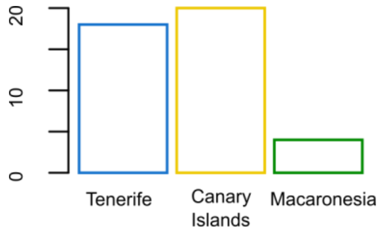


**Extended data Fig. 6 | Number of lineages present in Tenerife (18), Canary Islands (20) and Macaronesia (4).** The 21 major lineages (composed of 161 species or 34.7 % of all species included in the analysis) include a minimum three species per lineage. The 21 lineages are nested within the Macaronesia, Canary Islands and Tenerife.

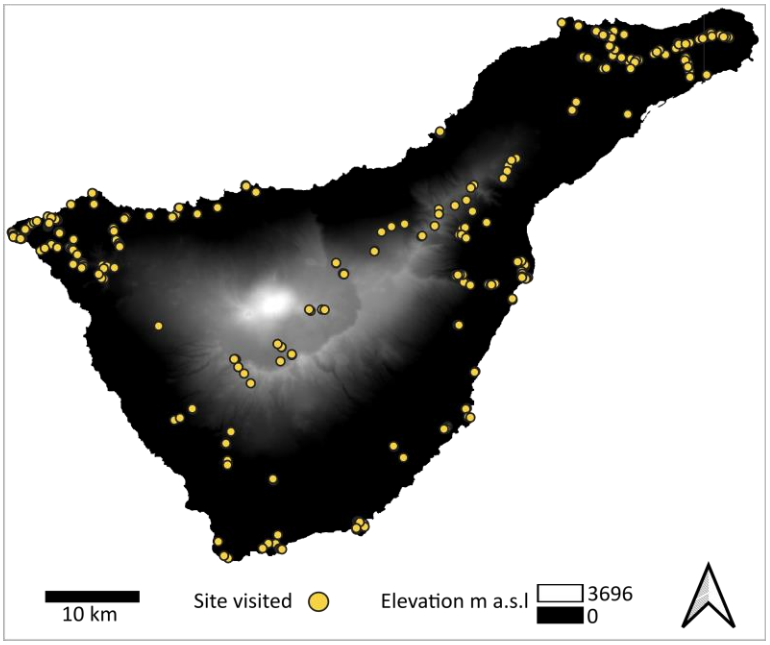


**Extended data Fig. 7 | Map of Tenerife with field sites visited**. In 2017 and 2018, we visited 500 different sites where plant materials were collected to measure plant functional traits.

**Extended data Table 1 | Validation of trait values imputation**. Out-of-Bag (OOB) scores are shown in units of the trait they correspond to. The lower the value the better the random forest algorithm performed. Note: Trait values predicted with phylogenetic information had, in almost all cases, lower OOB scores, except for LMA predictions.

| Imputed trait | OOB error with phylogenetically informed traits | OOB error with non-phylogenetically informed (naive) traits | Difference |
|---|---|---|---|
| Leaf area | 58079073.67 | 58213215.76 | -134142.1 |
| LDMC | 1018.36 | 1135 | -116.64 |
| LMA | 3050.18 | 2785.13 | 265.06 |
| Leaf th | 0.46 | 0.54 | -0.07 |
| Leaf N | 30.98 | 38.9 | -7.92 |
| Seed mass | 2184.05 | 2747.56 | -563.51 |
| Stem density | 0.02 | 0.02 | 0 |
| Height | 4.76 | 8.24 | -3.48 |

**Extended data Table 2. Assessment of functional contribution and functional originality of each radiated lineage to the island trait space**. We used Kruskal-Wallis tests with alpha level 0.05. Kruskal test comparison is done between lineage and non-'lineage'. Lineage group described using 'non-', e.g., non-Aeonium, includes all island seed plant native species except for species that belong to *Aeonium* lineage.

| Lineage group and non-lineage | Kruskal-Wallis Test Contribution | Kruskal-Wallis Test Originality | Lineage group and non-lineage | Kruskal-Wallis Test Contribution | Kruskal-Wallis Test Originality |
|---|---|---|---|---|---|
| *Aeonium* | a | a | *Limonium* | a | b |
| non-Aeonium | b | b | non-Limonium | a | a |
| *Argyranthemum* | b | b | *Lotus* | a | a |
| non-Argyranthemum | a | a | non-Lotus | a | a |
| *Aspagus* | a | a | *Micromeria* | a | a |
| non-Aspagus | a | a | non-Micromeria | a | a |
| *Bystropogon* | a | b | *Pericallis* | a | a |
| non-Bystropogon | a | a | non-Pericallis | a | a |
| *Cheirolophus* | b | b | *Pimpinella* | a | a |
| non-Cheirolophus | a | a | non-Pimpinella | a | a |
| *Convolvulus* | a | a | *Polycarpaea* | a | a |
| non-Convolvulus | a | a | non-Polycarpaea | b | b |
| *Crambe* | a | a | *Sideritis* | b | b |
| non-Crambe | a | a | non-Sideritis | a | a |
| *Descurainia* | a | a | *Sonchus* | b | a |
| non-Descurainia | a | a | non-Sonchus | a | a |
| *Echium* | a | a | *Teline* | a | a |
| non-Echium | a | a | non-Teline | a | a |
| *Euphorbia* | a | a | *Tolpis* | a | a |
| non-Euphorbia | a | a | non-Tolpis | a | a |
| *Helianthemum* | b | b | | | |
| non-Helianthemum | a | a | | | |
